# Genome assembly of *Firmina major*, an endangered savanna tree species endemic to China

**DOI:** 10.1101/2024.09.09.610897

**Authors:** Jing Yang, Rengang Zhang, Yongpeng Ma, Yuqian Ma, Weibang Sun

**Author notes:** corresponding author: Weibang Sun.

## Abstract

The tree species *Firmiana major* was once dominant in the savanna vegetation of the arid hot valleys of southwest China, but was considered extinct in the wild in 1998. After eight small populations were relocated by thorough investigations between 2018 and 2020, the species was subsequently recognized as a Plant Species of Extremely Small Populations (PSESP) in China in need of urgent rescue. Moreover, due to severe human disturbance, other species in the tropical woody genus *Firmiana* are also endangered, and the species in this genus have almost all been listed as second-class National Protected Wild Plants in China. In order to guide future research into the conservation of this group, we present here the high-quality genome assembly of *F. major*. This is the first genome assembly in the genus *Firmiana*, and is 1.4 Gb in size. The assembly consists of 1.18 Gb repetitive sequences, 37,673 annotated genes and 31,965 coding genes.

## Background & Summary

The woody genus *Firmiana* Marsili (Malvaceae) includes 18 species. Ten of these species have a natural distribution that includes the tropical and subtropical areas of China, and seven are endemic to this region. Most of the fossil records of *Firmiana* have been discovered in East Asia. A specimen has recently been found from the middle Eocene deposits of central Tibet, demonstrating that this genus occurred in that area^1^. *Firmiana* is thought to have radiated millions of years ago^2-4^ and was likely to have been very abundant at this time, however, the genus is now endangered throughout its modern distribution and even in its centre of diversity. In 2021, China listed all members within the genus (with the exception of *F. simplex*) as the second-class National Protected Wild Plants.

The target species *F. major* (Fig. 1a) was once considered extinct in the wild by the International Union for Conservation of Nature (IUCN) in 1998, but a small population was rediscovered in 2001. Between 2018 and 2020, we conducted a thorough field survey and found seven other populations of *F. major* in the arid-hot valleys along the Jinsha River, China^5,6^, where *F. major* is one of the few dominant tree species in the savanna vegetation. In a previous study^6^, we found that this species is threatened by habitat loss, and because the trees were subject to massive overharvesting and exploitation for their bark in previous centuries. We then initiated pilot scheme for the conservation of *F. major* including research on conservation genetics, propagation, in situ conservation, and reintroduction^7^. Moreover, the species was assessed as EN in the *China Red List of Biodiversity-Higher Plants Volume (2020)*, and has also been included on the List of Yunnan Protected Plant Species with Extremely Small Populations (PSESP), which is a plan for the deployment of conservation actions to save the most highly threatened species in China^8,9^. From 2022-2023, we finished sample collection and investigation of other *Firmiana* species in China and found that *Firmiana* species have strong tolerance of drought and barren soil and can survive in extremely harsh conditions. Though most populations of *Firmiana* species were severely disturbed by human activities, the surviving individuals are able to germinate and grow in rock crevices on cliffs, allowing them to avoid the impacts. Furthermore, morphological confusion in some populations hinted at possible hybridization between species, which might hinder the conservation of this genus.

**Figure 1.**
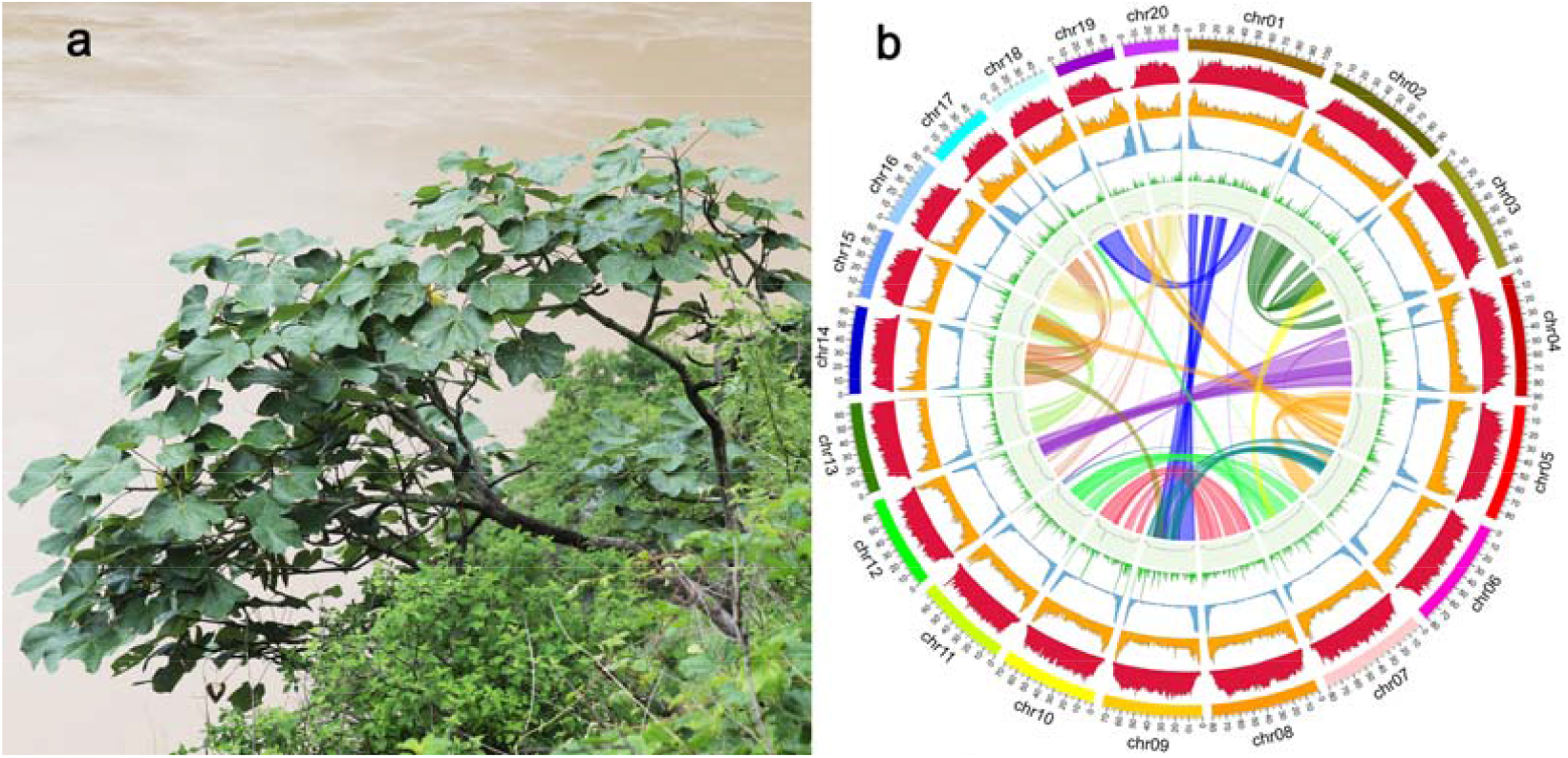
The tree of *F. major* (a) and the features of its genome assembly. The outmost circle represents all 20 chromosomes. The rings from outside to inside indicates class I TE density, class II TE density, protein-coding gene density, proportion of tandem repeat, GC density, genome rearrangement events of collinear blocks.

Here, we sequenced the genome of *F. major* and present the first high-quality chromosome-scale genome from the genus *Firmiana*. The genome of *F. major* is 1.4 Gb in size, accounting for 99.82 % of the 20 mapped chromosomes (Fig. 1b) and has contig and scaffold N50s of 56 Mb and 80 Mb, respectively. High LAI (17.74) and BUSCO (95.1 %) scores showed that the assembly had high sequence continuity and completeness, and a total of 1.18 Gb (83.3 %) repetitive sequences and 31,965 protein-coding genes were identified. This reference genome will greatly facilitate further studies in the conservation biology of *F. major* and other species in the genus of *Firmiana*, and will provide guidance for future conservation actions in this genus.

## Methods

### Sample preparation and Genome sequencing

Fresh *F. major* leaf material for genome library preparation and tissue samples (fruit, flower, leaf and twig) for transcriptome sequencing of *F. major* were collected from Dadong county in Lijiang City (LJ population, 27°09′ N, 100°26′ E)^6,7^, Yunnan Province, China, in 2021. Tissue samples were put into liquid nitrogen immediately after harvesting and preserved at −80 °C until extraction.

To ensure accurate assembly, we employed PacBio HiFi sequencing, Illumina sequencing and Hi-C sequencing using the PacBio Sequel II (Pacific Biosciences, USA) and Illumina Novaseq 6000 (Illumina, USA) plat forms. Total genomic DNA was extracted using a cetyltrimethyl ammonium bromide protocol^10^. A single-molecule real-time (SMRT) library was constructed and was used for whole-genome sequencing. The genomic DNA was sheared with Megaruptor 3 (Diagenode, USA) to an average size of 15-20 kb, then prepared using SMRTbell Template Prep Kit 1.0. The polymerase was bound using the Sequel II Binding Kit (v. 2.2) (Pacific Biosciences, USA), and the sequencing was performed on the PacBio Sequel II platform. Following the Illumina standard protocol, short-insert paired-end libraries were prepared and sequenced on the Illumina Novaseq 6000 platform with paired-end 150 read layout. Fastp^11^ (v. 0.19.3) was used to filter out the low-quality reads and adaptor sequences. The Hi-C library was prepared by cross-linking the chromosomal structure with formaldehyde, extraction of the genomic DNA, and digestion with the HindIII restriction enzyme, and was tagged with biotin-140dCTP and sheared into 300-600 bp fragments. Then, the Hi-C library was sequenced on the Illumina Novaseq 6000 platform. A total of 36 Gb PacBio HiFi reads (25 × coverage), 182 Gb Illumina short reads (130 × coverage) and 95 Gb Hi-C reads (68 × coverage) were obtained (Supplementary Table 1-3). RNA was extracted from tissue samples using TRIzol reagent (Invitrogen, USA). The PE RNA sequencing libraries were prepared using the NEBNEXT Ultra RNA Library Prep Kit for Illumina (New England Biolabs Inc.,USA) and 150 bp PE sequencing was performed on Illumina Hiseq X Ten platform, which generated 31 Gb reads (Supplementary Table 4).

### de novo assembly of the *F. major* genome

The PacBio HiFi reads were de novo assembled using the default parameters of Hifiasm^12^ (v. 0.13.0-R375), which generated a primary assembly at contig-level. The Hi-C data were then mapped to the contigs using Juicer^13^ (v. 1.5.6), and 3D-DNA^14^ (v. 180922) was utilized to scaffold a chromosome-level genome. Assembly errors were manually corrected with Juicerbox^15^ (v. 1.11.08). The 3D-DNA scaffolding pipeline was applied again to each chromosome to resolve the final scaffolds, followed by refinements in Juicebox, which generated chromosome scaffolds and un-anchored sequences. The genome assembly was then further improved by applying gap closing using TGS-GapClose^16^ (v. 1.1.1) with HiFi reads. For some chromosomes with incomplete telomeres, HiFi reads were mapped back to the chromosome with Minimap2^17^ (v. 2.24-r1122) and the reads aligned to chromosome ends were assembled to contigs using Hifiasm. By mapping the contigs back, the chromosome ends were extended. The chloroplast and mitochondrial genomes were assembled using the GetOrganelle toolkit^18^ (v. 1.7.5). The assembly was then polished using Nextpolish^19^ (v. 1.3.1) based on the Illumina short reads. Redundans^20^ (v. 0.13c) was used to eliminate the redundancy due to heterozygosity in the data. Thus, we obtained a high-quality genome assembly of *F. major*. The assembly is 1.4 Gb in size, and 99.82 % could be assigned to the 20 chromosomes. The genome had contig and scaffold N50s of 56 Mb and 80 Mb, respectively (Table 1). The chloroplast and mitochondrial genome sizes were 161,249 bp and 970,396 bp respectively.

**Table 1.**
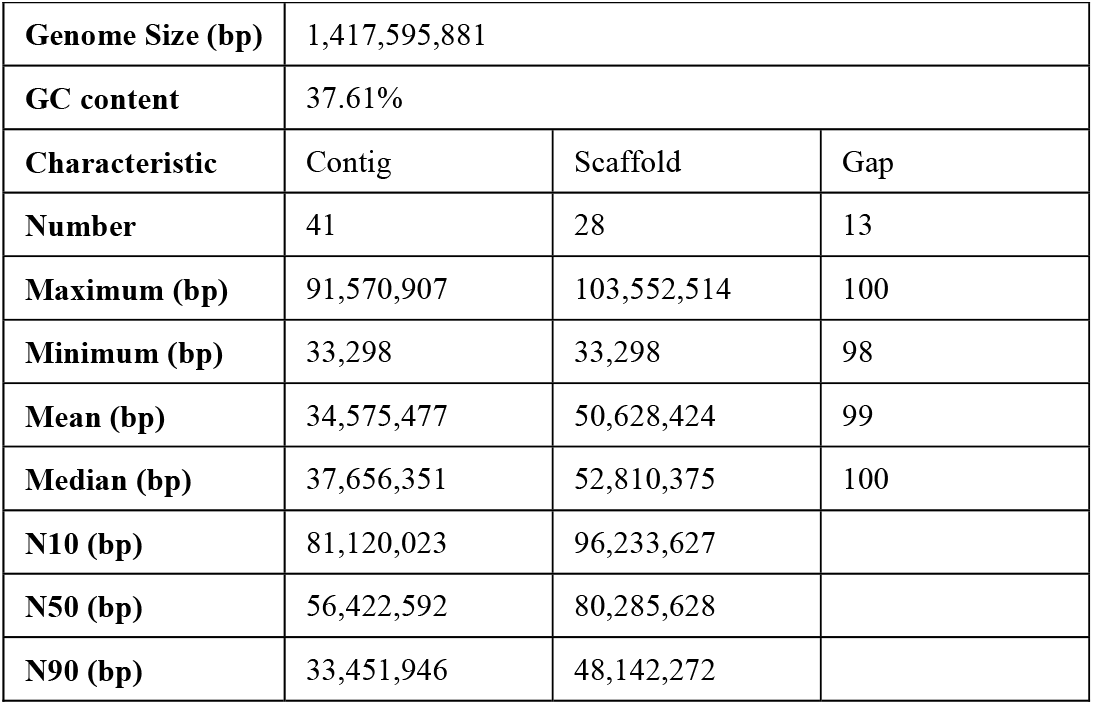
Statistics of the *Firmiana major* genome assembly.

|  |  |  |  |
| --- | --- | --- | --- |
| **Genome Size (bp)** | 1,417,595,881 |  |  |
| **GC content** | 37.61% |  |  |
| **Characteristic** | Contig | Scaffold | Gap |
| **Number** | 41 | 28 | 13 |
| **Maximum (bp)** | 91,570,907 | 103,552,514 | 100 |
| **Minimum (bp)** | 33,298 | 33,298 | 98 |
| **Mean (bp)** | 34,575,477 | 50,628,424 | 99 |
| **Median (bp)** | 37,656,351 | 52,810,375 | 100 |
| **N10 (bp)** | 81,120,023 | 96,233,627 |  |
| **N50 (bp)** | 56,422,592 | 80,285,628 |  |
| **N90 (bp)** | 33,451,946 | 48,142,272 |  |

### Annotation of repeats

A TE library of *F. major* was built using the de novo transposable elements annotator EDTA^21^ (v. 1.9.9; --sensitive 1 --anno 1). Against the library, RepeatMasker^22^ (v. 4.0.7) was employed to identify the repeat sequences within the assembled genome. A total of 2,289,617 repetitive sequences were screened, accounting for 83.3 % (1.18 Gb) of the genome. Long terminal repeats (LTRs) were found to be dominant (75.16 %, 1,06 Gb) (Table 2).

**Table 2.**
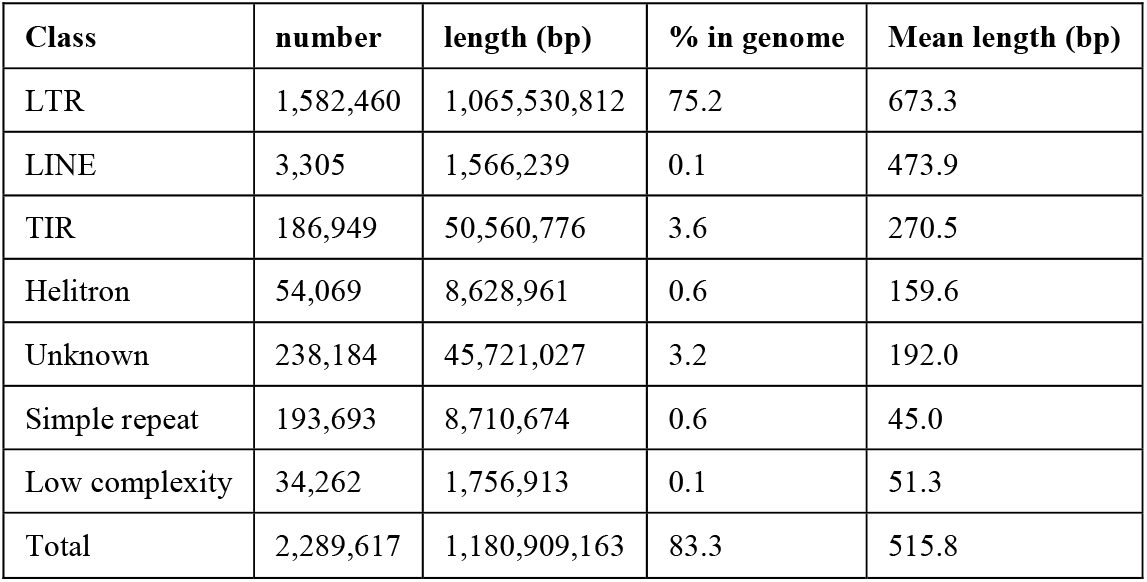
Repeat annotation statistics.

### Gene annotation

The transcriptome assembly was first prepared as RNA evidence for genome annotation. The RNA-seq reads were mapped to the genome assembly with HISAT2^23^ (v. 2.1.0). Genome-guided assembly was performed using Trinity (v. 2.0.6) and StringTie^24^ (v. 1.3.5). Trinity^25^ was used to generate the de novo assembly. The assembled transcripts were then clustered with CD-HIT^26^ (v. 4.6) and redundant transcripts were removed, which left 152,913 non-redundant transcripts (114.85 Mb, Supplementary Table 5). Based on these, the pipeline Program to Assemble Spliced Alignment (PASA)^27^ (v. 2.4.1) was used to annotate the structure of the genomic genes. By aligning the transcriptome to non-redundant protein sequences of *Durio zibethinus, Theobroma cacao, Gossypium raimondii, Hibiscus cannabinus, Corchorus capsularis, Aquilaria sinensis, Dipterocarpus turbinatus, Vitis vinifera* and *Arabidopsis thaliana*, 14,722 full-length genes were screened and used to train AUGUSTUS^28^ (v. 3.4.0) with five rounds of optimization.

The MAKER2 annotation pipeline^29^ was then used for genome annotation. Briefly, this included masking repeated regions with RepeatMasker (http://www.repeatmasker.org), ab initio prediction with AUGUSTUS, alignment of transcripts with BLASTN, alignment of protein evidence with BLASTX, refining alignments using Exonerate^30^ (v. 2.2.0), generating hints for AUGUSTUS, and yielding a high-quality gene model using AUGUSTUS by integrating predicted models. EVidenceModeler (EVM)^31^ (v. 1.1.1) was used to combine the gene models from MAKER2 and PASA to improve the accuracy of the final annotation results. This resulted in a consensus gene annotation, which was further modified in PASA by adding UTR sequences and alternative splicing. The process also removed overly short (< 50 aa) and abnormal gene annotations (e.g. those lacking start or stop codons, or containing internal stop codons). Moreover, tRNA, rRNA and other non-coding RNA were annotated using tRNAScan-SE^32^ (v. 1.3.1), barrnap (v. 0.9; https://github.com/tseemann/barrnap) and RfamScan^33^ (v. 14.2) respectively. After filtering out the redundancy, all annotation sets were integrated. A total of 37,673 annotated genes and 31,965 coding genes were acquired (Table 3).

**Table 3.** Gene annotation statistics.

| Feature | All genes |  |  |  |  | Coding genes |  |  |  |  |
| --- | --- | --- | --- | --- | --- | --- | --- | --- | --- | --- |
|  | number | min. | max. | median | mean | number | min. | max. | median | mean |
| gene | 37,673 | 53 | 272,505 | 2,240 | 3,346.6 | 31,965 | 153 | 272,505 | 2,748 | 3,914.6 |
| transcript | 61,430 | 53 | 19,336 | 1,659 | 1,806.3 | 55,722 | 153 | 19,336 | 1,779 | 1,974.4 |
| CDS | 55,722 | 153 | 18,999 | 1,086 | 1,312.5 | 55,722 | 153 | 18,999 | 1,086 | 1,312.5 |
| exon | 368,519 | 3 | 10,426 | 151 | 301.1 | 362,782 | 3 | 10,426 | 154 | 303.3 |
| intron | 307,089 | 4 | 266,343 | 186 | 478.1 | 307,060 | 21 | 266,343 | 186 | 478.2 |
| exons/transcript | 61,430 | 1 | 77 | 5 | 6 | 55,722 | 1 | 77 | 5 | 6.5 |

The functions of protein-coding genes were predicted using three methods. First, the eggNOG Orthologous Groups database and eggNOG-mapper^34^ (v. 2.1.6) were used for the aligning of homologous genes and for functional annotation. Then the Swiss-Prot, TrEMBL and NCBI-nr databases were used for alignment and annotation of protein sequence using DIAMOND^35^ (v. 0.9.24; Identity > 30 % and E-value < 1e-5). Thirdly, InterProScan^36^ (v. 5.27-66.0) was used for assignment based on domain conservation, and comparison with InterPro’s signatures provided by databases including PRINTS, Pfam, SMART, PANTHER, CDD, and others. Finally, 31,486 genes were functionally annotated (Table 4).

**Table 4.**
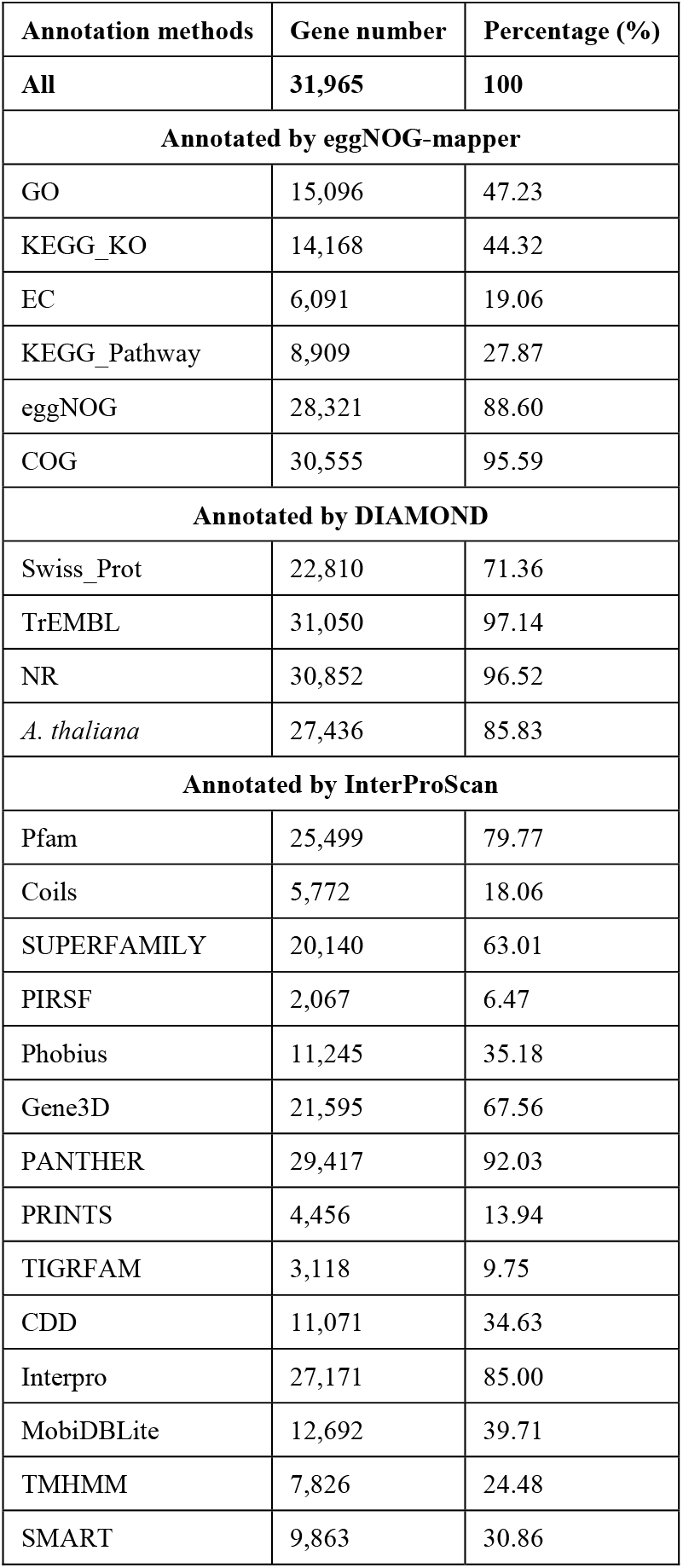
Function annotation of the predicted protein-coding genes.

| Annotation methods | Gene number | Percentage (%) |
| --- | --- | --- |
| All | 31,965 | 100 |
| Annotated by eggNOG-mapper |  |  |
| GO | 15,096 | 47.23 |
| KEGG_KO | 14,168 | 44.32 |
| EC | 6,091 | 19.06 |
| KEGG_Pathway | 8,909 | 27.87 |
| eggNOG | 28,321 | 88.60 |
| COG | 30,555 | 95.59 |
| Annotated by DIAMOND |  |  |
| Swiss_Prot | 22,810 | 71.36 |
| TrEMBL | 31,050 | 97.14 |
| NR | 30,852 | 96.52 |
| *A. thaliana* | 27,436 | 85.83 |
| Annotated by InterProScan |  |  |
| Pfam | 25,499 | 79.77 |
| Coils | 5,772 | 18.06 |
| SUPERFAMILY | 20,140 | 63.01 |
| PIRSF | 2,067 | 6.47 |
| Phobius | 11,245 | 35.18 |
| Gene3D | 21,595 | 67.56 |
| PANTHER | 29,417 | 92.03 |
| PRINTS | 4,456 | 13.94 |
| TIGRFAM | 3,118 | 9.75 |
| CDD | 11,071 | 34.63 |
| Interpro | 27,171 | 85.00 |
| MobiDBLite | 12,692 | 39.71 |
| TMHMM | 7,826 | 24.48 |
| SMART | 9,863 | 30.86 |

### Data Records

The raw sequencing data from the PacBio HiFi reads, Illumina short reads, Hi-C reads and transcriptomic sequencing were deposited in the NCBI Sequence Read Archive database under the BioProject accession number PRJNA1007875. The genome assembly was deposited in GenBank under the accession number JAVKOW000000000^37^. The data mentioned were also deposited in the Genome Warehouse of the National Genomics Data Center, Beijing Institute of Genomics, Chinese Academy of Sciences / China National Center for Bioinformation, under the accession number GWHDTXC00000000^38, 39^.

### Technical Validation

Based on the LTR (long terminal repeat) assembly index (LAI)^40^, the genome assembly was evaluated with a score of 17.74, which indicated high contiguity (categorized as reference level when 10 ≤ LAI ≤ 20). The Illumina short reads, HiFi reads and the RNA sequencing reads were mapped to the assembly using BWA^41^ (v. 0.7.17), Minimap2^17^ (v. 2.24) and HISAT2^23^ (v. 2.1.0) respectively, and the quality was also validated by the high mapping rates achieved (Table 5).

**Table 5.** Assessment of genome coverage rate.

| Data set | Reads mapped | Properly paired | Bases mapped | $\geq 1\times$ | $\geq 5\times$ | $\geq 10\times$ | $\geq 20\times$ |
| --- | --- | --- | --- | --- | --- | --- | --- |
| Next generation | 99.9% | 96.9% | 99.9% | 99.8% | 99.2% | 98.6% | 97.4% |
| HiFi | 99.8% | - | 99.8% | 99.99% | 99.9% | 98.5% | 75.9% |
| RNA-Seq | 93.4% | 91.3% | 93.4% | 8.9% | 6.2% | 5.1% | 4.1% |

Furthermore, the quality of the final assembly was assessed using BUSCO^42^ (v. 5.3.2), which resulted in a high completeness score of 95.1 % (Table 6). By plotting the distribution of coverage depth across the whole genome against the sequencing data (Figure 2), we found that duplicated genes had the same depth distribution as single-copy genes and also matched the Poisson distribution. No obvious heterozygous peak was observed, and, combining the above data, we concluded that the assembly had no redundancy. SAMtools/BCFtools (v. 1.9) was employed for variant calling, which found a heterozygosity rate of 0.24 % and an error rate of 0.00044 %. In the guanine-cytosine (GC) depth analysis, the genome was found to have a mean GC content of 37.61

**Table 6.**
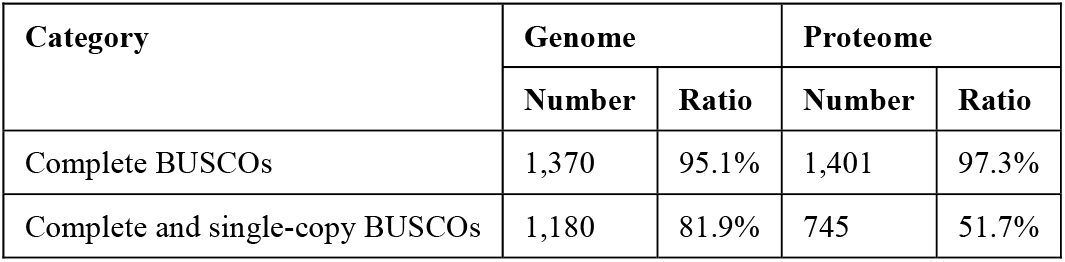
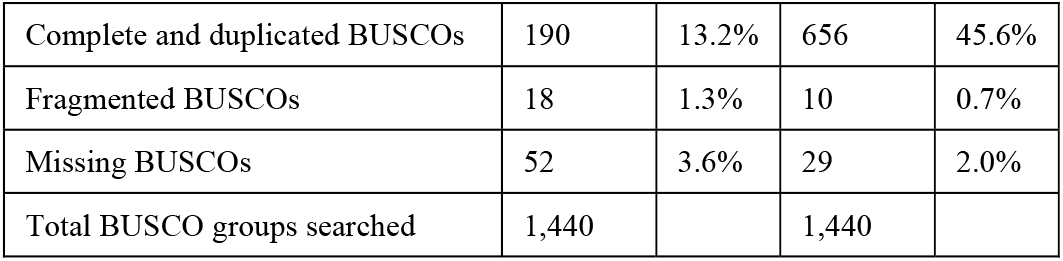
Assessment of BUSCO.

| Category | Genome |  | Proteome |  |
| --- | --- | --- | --- | --- |
|  | Number | Ratio | Number | Ratio |
| Complete BUSCOs | 1,370 | 95.1% | 1,401 | 97.3% |
| Complete and single-copy BUSCOs | 1,180 | 81.9% | 745 | 51.7% |
| Complete and duplicated BUSCOs | 190 | 13.2% | 656 | 45.6% |
| Fragmented BUSCOs | 18 | 1.3% | 10 | 0.7% |
| Missing BUSCOs | 52 | 3.6% | 29 | 2.0% |
| Total BUSCO groups searched | 1,440 |  | 1,440 |  |

**Figure 2.**
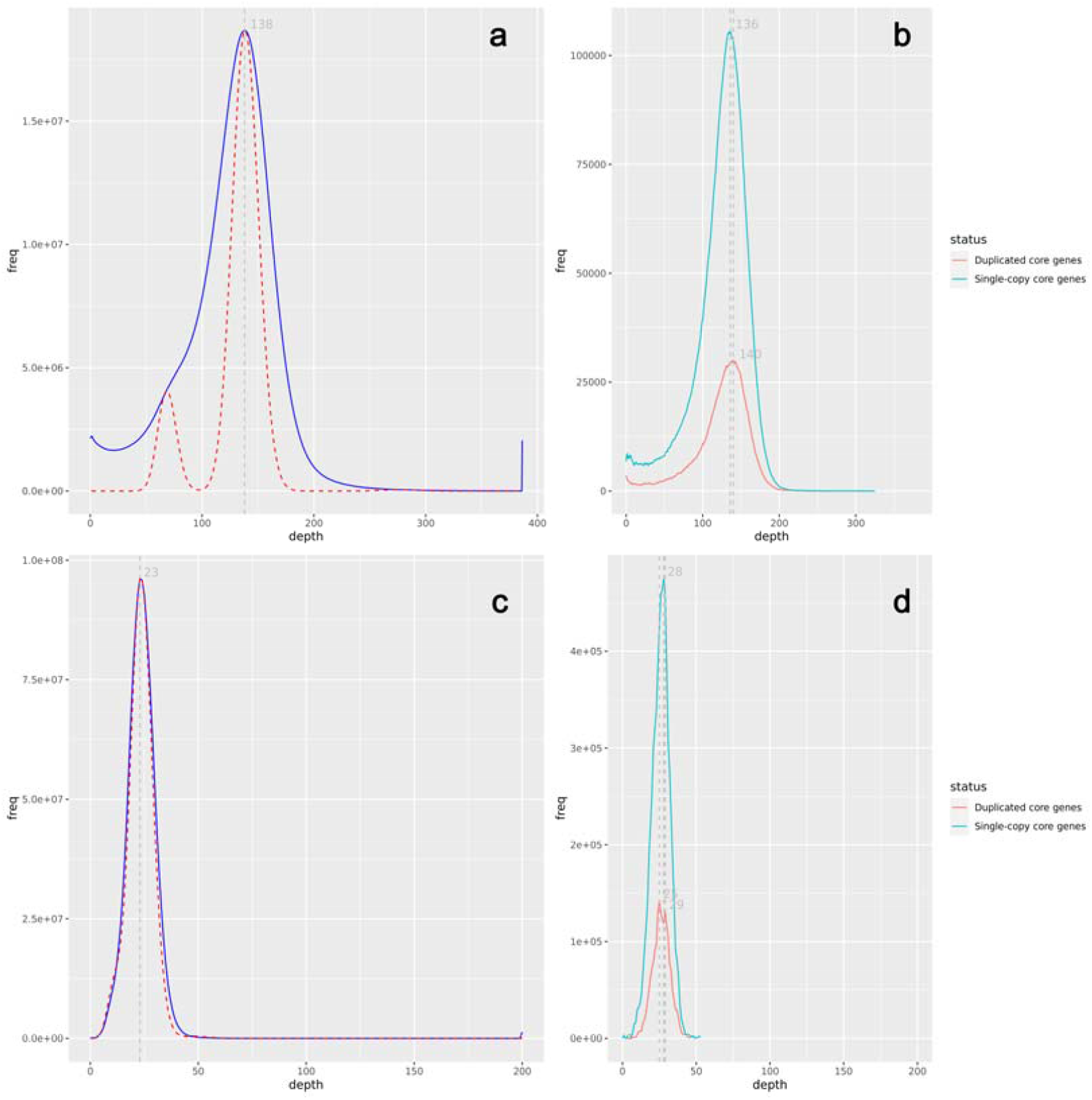
Coverage depth of genomic assembly (a) and BUSCO core gene set (b) using Illumina sequencing data. Coverage depth of genomic assembly (c) and BUSCO core gene set (d) using HiFi sequencing data.

%, and a GC bias was found in the Illumina sequencing data (Figue 3). After Juicer was used to map the Hi-C data to the genome, the 20 chromosome scaffolds showed no obvious assembly errors (Figue 4). The repetitive sequences were mapped to the genome, demonstrating that most telomere sequences were complete and 18-5.8-28S rDNA arrays were detected on Chr13. In conclusion, these assessments indicate that the genome assembly was complete and of high quality. The quality of gene annotation was also evaluated using BUSCO (Table 6). The complete core gene accounted for a proportion of 97.3%. Only 0.7% genes had fragmented matches and 2.0% were missing.

**Figure 3.**
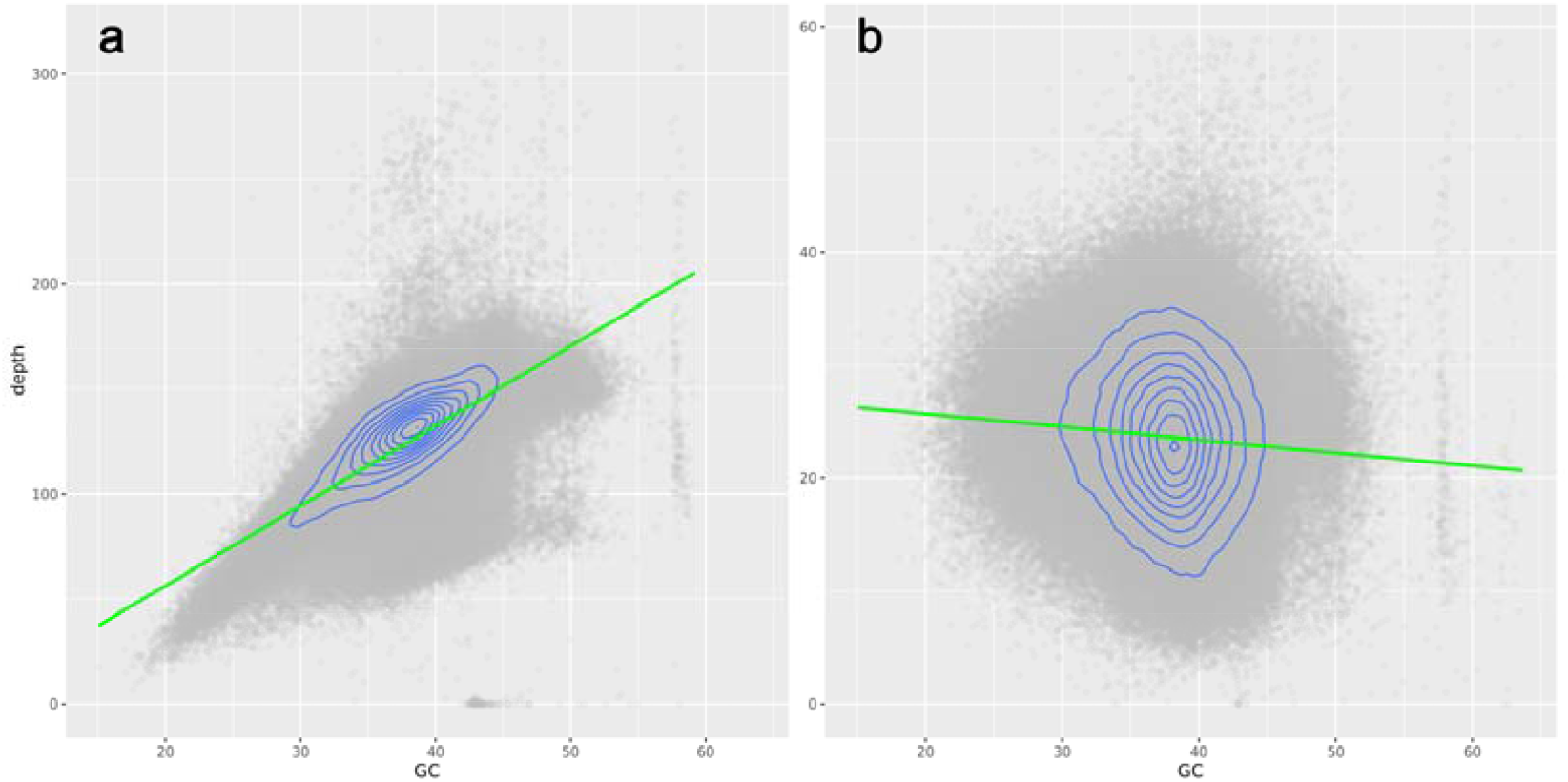
Coverage depth of Illumina sequencing data (a) and HiFi sequencing data (b) at different GC contents.

**Figure 4.**
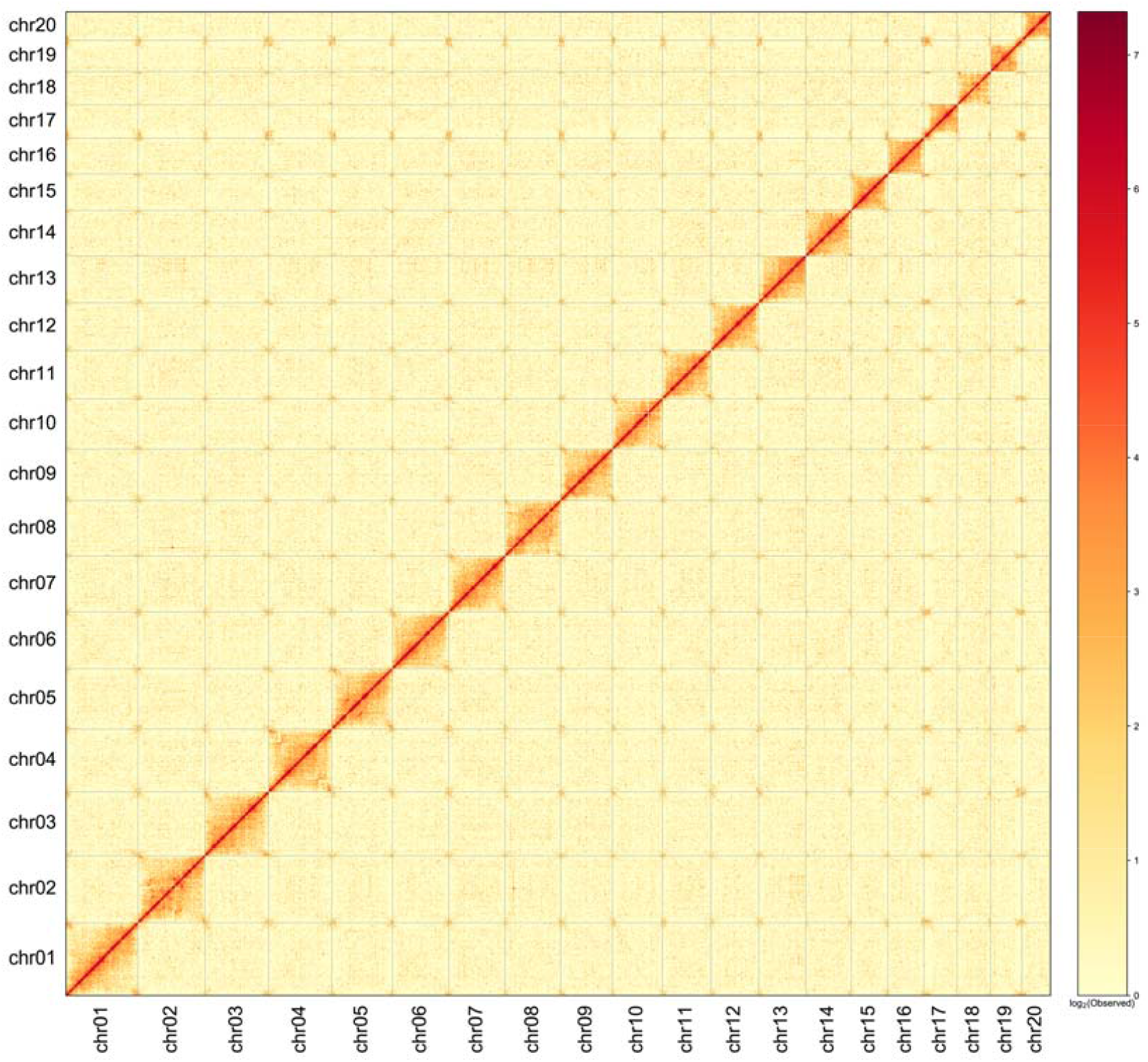
Hi-C heatmap demonstrated 20 unambiguous chromosome scaffolds with no obvious chromosome assembly error.

## Code Availability

The software and pipelines mentioned in the paper were executed following published manuals and protocols.

## Acknowledgements

The research was supported by the Second Tibetan Plateau Scientific Expedition and Research (STEP) Program (2019QZKK0502), the Science and Technology Basic Resources Investigation Program of China (2017FY100100), the National Natural Science Foundation of China (32371743), the Natural Science Foundation of Yunnan Province (202201AS070049), the CAS “Light of West China” Program (to Jing Yang and Yongpeng Ma) and the Ten Thousand Talent Program of Yunnan Province (YNWR-QNBJ-2019-127 and YNWR-QNBJ-2018-174).

## Author contributions

Weibang Sun, Yongpeng Ma and Jing Yang conceived and designed the study. Yuqian Ma collected and prepared the samples. Rengang Zhang performed the analyses. Jing Yang drafted the manuscript. Rengang Zhang and Weibang Sun modified the manuscript. All authors have read, revised, and approved the final manuscript for submission.

## Competing interests

The authors declare that no competing interests involved in this research.

## Notes

### Competing Interest Statement

The authors have declared no competing interest.

## References

1. Del Rio, C. et al. Fruits of Firmiana and Craigia (Malvaceae) from the Eocene of the Central Tibetan Plateau with emphasis on biogeographic history. Journal of Systematics and Evolution 60, 1440–1452 (2022).

2. Srivastava, G. & Mehrotra, R. C. Further contribution to the low latitude leaf assemblage from the late Oligocene sediments of Assam and its phytogeographical significance. Journal of Earth System Science 122, 1341–1357 (2013).

3. Xie, S., Manchester, S. R., Liu, K., Wang, Y. & Shao, Y. Firmiana (Malvaceae: Sterculioideae) fruits from the Upper Miocene of Yunnan, Southwest China. Geobios 47, 271–279 (2014).

4. Jia, L. et al. Fossil fruits of Firmiana and Tilia from the middle Miocene of South Korea and the efficacy of the Bering land bridge for the migration of mesothermal plants. Plant Diversity 43, 480–491 (2021).

5. Yang, J., Chen, G. & Sun, W. Conserving Firmiana major, a tree species endemic to China. Oryx 52, 211–211 (2018).

6. Li, C. et al. Population structure and regeneration dynamics of Firmiana major, a dominant but endangered tree species. Forest Ecology and Management 462, 117993 (2020).

7. Ma, Y. et al. Conservation genetics of Firmiana major, a threatened tree species with potential for afforestation of hot, arid climates. Global Ecology and Conservation 36, e02136 (2022).

8. Yang, J. et al. China’s conservation program on Plant Species with Extremely Small Populations (PSESP): Progress and perspectives. Biological Conservation 244, 108535 (2020).

9. Sun, W. List of Yunnan protected plant species with extremely small populations. (Yunnan Science and Technology Press, 2021).

10. Porebski, S., Bailey, L. G. & Baum, B. R. Modification of a CTAB DNA extraction protocol for plants containing high polysaccharide and polyphenol components. Plant Molecular Biology Reporter 15, 8–15 (1997).

11. Chen, S., Zhou, Y., Chen, Y. & Gu, J. fastp: an ultra-fast all-in-one FASTQ preprocessor. Bioinformatics 34, 884–890 (2018).

12. Cheng, H., Concepcion, G. T., Feng, X., Zhang, H. & Li, H. Haplotype-resolved de novo assembly using phased assembly graphs with hifiasm. Nature Methods 18, 170–+ (2021).

13. Durand, N. C. et al. Juicer provides a one-click system for analyzing loop-resolution Hi-C experiments. Cell Systems 3, 95–98 (2016).

14. Dudchenko, O. et al. De novo assembly of the Aedes aegypti genome using Hi-C yields chromosome-length scaffolds. Science 356, 92–95 (2017).

15. Durand, N. C. et al. Juicebox provides a visualization system for Hi-C contact maps with unlimited zoom. Cell Systems 3, 99–101 (2016).

16. Xu, M. et al. TGS-GapCloser: a fast and accurate gap closer for large genomes with low coverage of error-prone long reads. GigaScience 9, giaa094 (2020).

17. Li, H. Minimap2: pairwise alignment for nucleotide sequences. Bioinformatics 34, 3094–3100 (2018).

18. Jin, J. et al. GetOrganelle: a fast and versatile toolkit for accurate de novo assembly of organelle genomes. Genome Biology 21, 241 (2020).

19. Hu, J., Fan, J., Sun, Z. & Liu, S. NextPolish: a fast and efficient genome polishing tool for long-read assembly. Bioinformatics 36, 2253–2255 (2020).

20. Pryszcz, L. P. & Gabaldón, T. Redundans: an assembly pipeline for highly heterozygous genomes. Nucleic acids research 44, e113–e113 (2016).

21. Ou, S. et al. Benchmarking transposable element annotation methods for creation of a streamlined, comprehensive pipeline. Genome Biology 20, 275 (2019).

22. Tarailo-Graovac, M. & Chen, N. Using RepeatMasker to identify repetitive elements in genomic sequences. Current protocols in bioinformatics Chapter 4, 4.10.11-14.10.14 (2009).

23. Kim, D., Langmead, B. & Salzberg, S. L. HISAT: a fast spliced aligner with low memory requirements. Nature Methods 12, 357–360 (2015).

24. Pertea, M. et al. StringTie enables improved reconstruction of a transcriptome from RNA-seq reads. Nature Biotechnology 33, 290–295 (2015).

25. Grabherr, M. G. et al. Full-length transcriptome assembly from RNA-Seq data without a reference genome. Nature Biotechnology 29, 644–652 (2011).

26. Li, W. & Godzik, A. Cd-hit: a fast program for clustering and comparing large sets of protein or nucleotide sequences. Bioinformatics 22, 1658–1659 (2006).

27. Haas, B. J. et al. Improving the Arabidopsis genome annotation using maximal transcript alignment assemblies. Nucleic Acids Research 31, 5654–5666 (2003).

28. Stanke, M., Diekhans, M., Baertsch, R. & Haussler, D. Using native and syntenically mapped cDNA alignments to improve de novo gene finding. Bioinformatics 24, 637–644 (2008).

29. Holt, C. & Yandell, M. MAKER2: an annotation pipeline and genome-database management tool for second-generation genome projects. BMC Bioinformatics 12, 491 (2011).

30. Slater, G. S. C. & Birney, E. Automated generation of heuristics for biological sequence comparison. BMC Bioinformatics 6, 31 (2005).

31. Haas, B. J. et al. Automated eukaryotic gene structure annotation using EVidenceModeler and the Program to Assemble Spliced Alignments. Genome Biology 9, R7 (2008).

32. Lowe, T. M. & Eddy, S. R. tRNAscan-SE: A program for improved detection of transfer RNA genes in genomic sequence. Nucleic Acids Research 25, 955–964 (1997).

33. Kalvari, I. et al. Non-coding RNA analysis using the Rfam Database. Curr Protoc Bioinformatics 62, e51 (2018).

34. Huerta-Cepas, J. et al. Fast genome-wide functional annotation through orthology assignment by eggNOG-Mapper. Molecular Biology and Evolution 34, 2115–2122 (2017).

35. Buchfink, B., Xie, C. & Huson, D. H. Fast and sensitive protein alignment using DIAMOND. Nature Methods 12, 59–60 (2015).

36. Jones, P. et al. InterProScan 5: genome-scale protein function classification. Bioinformatics30, 1236–1240 (2014).

37. NCBI Assembly https://identifiers.org/ncbi/insdc.gca:JAVKOW000000000 (2023).

38. Genome Warehouse: A Public Repository Housing Genome-scale Data. Genom. Proteom.Bioinfo. 19, 584–589 (2021).

39. Database Resources of the National Genomics Data Center, China National Center for Bioinformation in 2023. Nucleic Acids Res 2023, 51(D1):D18–D28.

40. Ou, S., Chen, J. & Jiang, N. Assessing genome assembly quality using the LTR Assembly Index (LAI). Nucleic Acids Research 46, e126–e126 (2018).

41. Li, H. Aligning sequence reads, clone sequences and assembly contigs with BWA-MEM.arXiv 1303, 3997 (2013).

42. Simão, F. A., Waterhouse, R. M., Ioannidis, P., Kriventseva, E. V. & Zdobnov, E. M. BUSCO: assessing genome assembly and annotation completeness with single-copy orthologs. Bioinformatics 31, 3210–3212 (2015).

